# Validation of RT-qPCR primers and probes for new and old variants of SARS-CoV-2 in a world scale

**DOI:** 10.1101/2024.03.19.585194

**Authors:** Alderrosy Fragoso Rodrigues Almeida, Weriskiney Araújo Almeida, Wvelton Mendes Pereira, Renato de Santana Silva, Larissa Paola Rodrigues Venancio, Mary Hellen Fabres-Klein, Jonilson Berlink Lima, Raphael Contelli Klein, Théo Araújo-Santos

## Abstract

**Introduction:** The demand for molecular diagnosis of pathogens has surged dramatically since the onset of the COVID-19 pandemic. In this context, different diagnostic tests have been developed to identify SARS-CoV-2 in patient samples. The emergence of new variants of SARS-CoV-2 raises questions about whether the molecular tests available for diagnosis continue to be effective in detecting the virus in biological samples.

**Objective:** This study analyzed the viability of molecular targets directed to N, E and RdRp genes available against the new variants of SARS-CoV-2.

**Methodology:** For this, we used bioinformatics tools to analyze SARS-CoV-2 genomic data of different variants deposited in GSAID and NCBI virus genomic databases to assess the accuracy of molecular tests available for the diagnosis of COVID-19. We also developed software for analyzing mutation frequencies in different molecular targets from the mutation database.

**Results:** Mutation frequency analysis revealed a high rate of mutations in the N, E and RdRp genes and targets, although the target regions were more conserved. Only three SNPs were recurrent in the sequences of the variants identified in different continents and all in different targets. On the other hand, the registered mutations are not consistent and do not appear frequently in isolates of the same variant in all regions of the world.

**Conclusion:** Our data suggest that the molecular targets designed for the first SARS-CoV-2 variants remain valid for the identification of new virus variants despite the large number of identified haplotypes. However, false negative test failures can be identified by using more than one molecular target for the same sample. Genomic regions that are under evolutive selective pressure should be avoided in the use of the diagnostic, once the emergence of new variants may affect the efficiency of molecular testing on a global scale.

## INTRODUCTION

The demands of molecular diagnosis for coronavirus disease 2019 (COVID-19) have democratized the use of the quantitative real-time polymerase chain reaction (RT-qPCR) technique and exposed its limitations. Different RT-qPCR kits used to identify severe acute respiratory syndrome coronavirus 2 (SARS-CoV-2) were made available around the world, targeting different regions of the viral genome [1–4]. However, due to the specific characteristics of RNA viruses like the increasing and rapid number of mutations and consequently emergence of viral variants may constitute an obstacle to the diagnosis of SARS-CoV2, leading to invalidation of molecular targets [5]. The variants were categorized by the WHO into two groups called Variants of Interest (VOIs) and Variants of Concern (VOCs). VOIs are variants that present mutations that interfere with transmission, severity, detection and sensitivity to medicines, as well as causing significant community transmission, or an increase in cases in some countries, presenting prevalence during or over time, or even those that may pose risks to the population [6]. VOCs, in turn, comprise the characteristics of VOIs plus others, such as increased transmissibility, virulence or clinical presentation of the disease, or decreased effectiveness of social and public health measures (https://www.who.int/publications/m/item/historical-working-definitions-and-primary-actions-for-sars-cov-2-variants, accessed on 18 December, 2023).

In a recently published study, our group demonstrated that it is possible to identify SARS-CoV-2 haplotypes using bioinformatics techniques and predict the viability of molecular targets for the diagnosis of COVID-19 in commercial tests [1].

After three years of the COVID-19 pandemic, Brazil is still among the main countries affected by the disease and, despite constant financial cuts in scientific areas, it continues to be among the countries that deposit the most sequences in SARS-CoV2 genomic databases [7]. In the context of genomic surveillance of the virus, groups such as GSAID (Global Data Science Initiative) have been contributing to the curation of available sequences of SARS-CoV-2 variants, allowing researchers from around the world to verify the viability of molecular tests available around the world [7,8].

In this work, we verified the validity of the molecular targets (N, E, RdRp) used to identify the SARS-CoV-2 virus, by analyzing the frequency of single base polymorphisms deposited in databases. In addition, we purpose an easy workflow to identify possible pitfalls in the molecular tests to detection of SARS-CoV-2 new variants.

## METHODS

### Selection of primers and probes used to identify SARS-CoV-2

The sequences identification of primers and probes used in molecular diagnostic methods was conducted through a literature review using the PUBMED database with the following descriptors: SARS-CoV-2, RT-PCR, RT-qPCR, qPCR, Real time PCR, molecular diagnosis, COVID-19, Primer. Only articles that presented primer sequences and probes for detecting the virus were included in the study. The publication period of the works covered the years 2020 to 2023. The protocols selected are summarized in the Table 1.

**Table 1.**
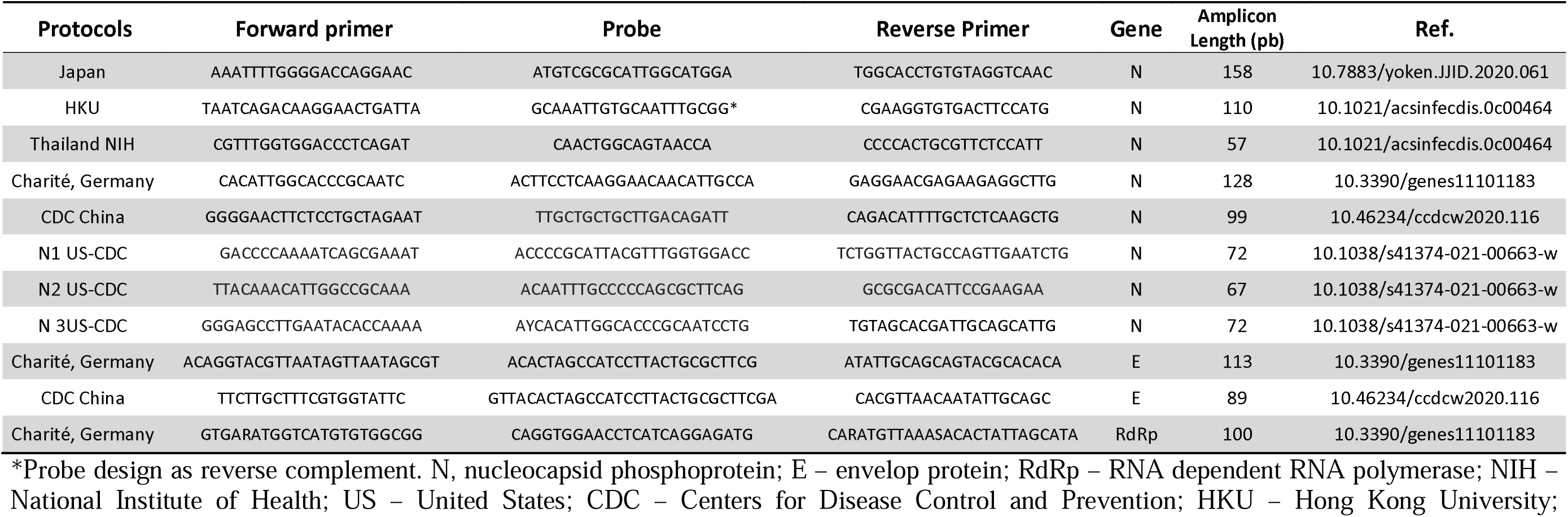
The most used primer and probe sets for SARS-CoV-2 diagnostic by RT-qPCR.

### SARS-CoV-2 variant sequences

At least ten sequences representing each variant of concern (VOC), P.1, B.1.1.7, B.1.617.2, B.1.351, B.1.1.529, were collected from the GSAID website (https://gisaid. org/). The regions of interest were compared using the MEGA 11 software [9] (Supplemental Material). The selection of sequences in GSAID was as follows, in order of priority: (i) Complete sequences and/or sequences with high coverage; (ii) Representatives from different continents; (iii) Earlier deposit date; (iv) The status of identified patients.

### Software Target Mutations® to identification of SNPs in the primers for RT-qPCR

Single nucleotide polymorphisms in the E and NP gene were identified by using available data in the National Center for Biotechnology Information https://www.ncbi.nlm.nih.gov/labs/virus/vssi/#/scov2_snp; accessed February 27, 2023). Next, we measured the absolute and relative frequencies of the SNPs present in the different primer and probe sets used in the RT-qPCR to detect SARS-CoV-2 using as reference genome of SARS-CoV-2 (NC_045512.2). To measure the mutation frequency in the SARS-CoV-2 genes, primers and probe set of the RT-qPCR protocols, we develop a software named *“Target Mutations® 1.0”* (Barreiras, BA, Brazil) to quantify and provide quality of the SNPs (INPI register number # BR512023002633-6). The software was constructed using HTML, Java Script and CSS program languages (Software and Supplemental Material). Briefly, the software uses as entry the primer and/or probe nucleotide sequences and then identify the mutations in the onboard Database of the mutations (Software script and Database). Next, the *Target Mutations®* provides absolute and relative frequencies of the synonymous and non-synonymous mutations of the hole gene and/or primer and probe analyzed and download the results as a comma-separated values file extension (.csv) (Software in Supplemental Material).

### Long oligo design to RT-qPCR validation

Long oligos were designed to mimetize SNPs found in new variants in regions of the annealing primers or probes of E or RdRp gene targets. Two SNPs present in the annealing regions most frequently in new SARS-CoV-2 variants were evaluated. For PCR tests, serial dilutions of 1:10 of the long oligos were used, starting at 1×10^6^ template molecules per 1 µL of reaction. Long oligo sequences were E: 5’- ATAGGTACGTTAATAGTTAATAGCGTACTTCTTTTTCTTGCTTTCGTGGTATTCTTGCTAGTTACACTAGCCATCCTTACTGCGCTTCGATTGTGTGCGTACTGCTGCAATAT-3’ and amplicon RdRp: 5′- GTGAAATGGTCATGTGTGGCAGTTCACTATATGTTAAACCAGGTGGAACCTCATCAGGAGATGCCACAACTGCTTATGCTAATAGTGTTTTTAACATTTG-3’.

### RT-qPCR

Reverse transcription quantitative PCR analysis was performed on QuantStudio 5 Real-Time PCR system (Thermo Scientific, USA) using the primer set from E or RdRp Charité (IDT Coralville, IA). The PCR reaction mixture consisted of Kappa Probe Fast qPCR Master Mix (2X) Kit (Sigma-Aldrich), 0.75 μL of primers and 2.5 μL of RNA in a final volume of 10 μL reaction. Cycling conditions were 42 °C for 5 min and 95 °C for 3 minutes, followed by 45 cycles at 95 °C for 5 seconds and 60 °C for 1 minute. The primers and concentrations used in the experiment were as follows: 400 ηM E: Forward: 5′- ACAGGTACGTTAATAGTTAATAGCGT-3′; 400 ηM E: Reverse: 5′- ATATTGCAGCAGTACGCACACA-3′; 200 ηM E: Probe FAM-ACACTAGCCATCCTTACTGCGCTTCG-NFQ-MGB; 600 ηM RdRp: Forward: 5′- GTGARATGGTCATGTGTGGCGG-3′; 800 ηM RdRp: Reverse: 5′- CARATGTTAAASACACTATTAGCATA-3′ and 100 ηM RdRp: Probe FAM-CAGGTGGAACCTCATCAGGAGATG-NFQ-MGB [10]. Alternatively, Allplex^TM^ 2019-nCoV assay (Seegene, South Korea) was used to identified long oligos described above, since this assay use the same primer and probe set of E and RdRp Charité protocol [10].

### Statistical Analysis

Absolute and relative frequencies were quantified using Target Mutations*®* (Barreiras, BA, Brazil) and the data were used to construct the representative graphs using GraphPad-Prism 5.0 software (GraphPad Software, San Diego, CA-USA).

## RESULTS

### Mutation frequency in the RT-qPCR molecular targets used to SARS-CoV-2 identification

Many RT-qPCR protocols were previously stablished to detection of SARS-CoV-2 in samples of infected patients. Herein, we verified the variability in the sets of primers and probes used in the protocols during the first wave of the COVID-19 pandemic (Table 1). Presence of SNPs in the amplicons regions has been reported as invalidating several RT-qPCR protocols [5]. We identified the mutation frequencies in the amplicon regions of the E and N gene targets, which were extensive used around the world to SARS-CoV-2 detection (Figure 1A). Consortium initiative to identify SNPs in the SARS-CoV-2 genomes make disponible more than twenty-six thousand SNPs in whole virus sequence (Database in Supplemental Material). We worked in data mining, and we identified more mutations out of amplicon regions independently of gene analyzed (Figure 1A). Protein of envelop gene presented higher frequency of mutations than N gene, which it can be explained since E gene has a very short sequence (228 nucleotides) compared to N gene (1260 nucleotides) (Figure 1A). Genomic mutations can be considered important if they confer some evolutionary advantage to the haplotypes. In this sense, non-synonymous (NS) mutations that are related to modification in the aminoacids residues in the protein coded are more relevant and possibly conserved [11]. We found more NS mutations in the amplicon regions of the E gene compared N gene (Figure 1B). Despite this, the data indicate that both the E and N genes present mutations throughout their entire sequence.

**FIGURE 1.**
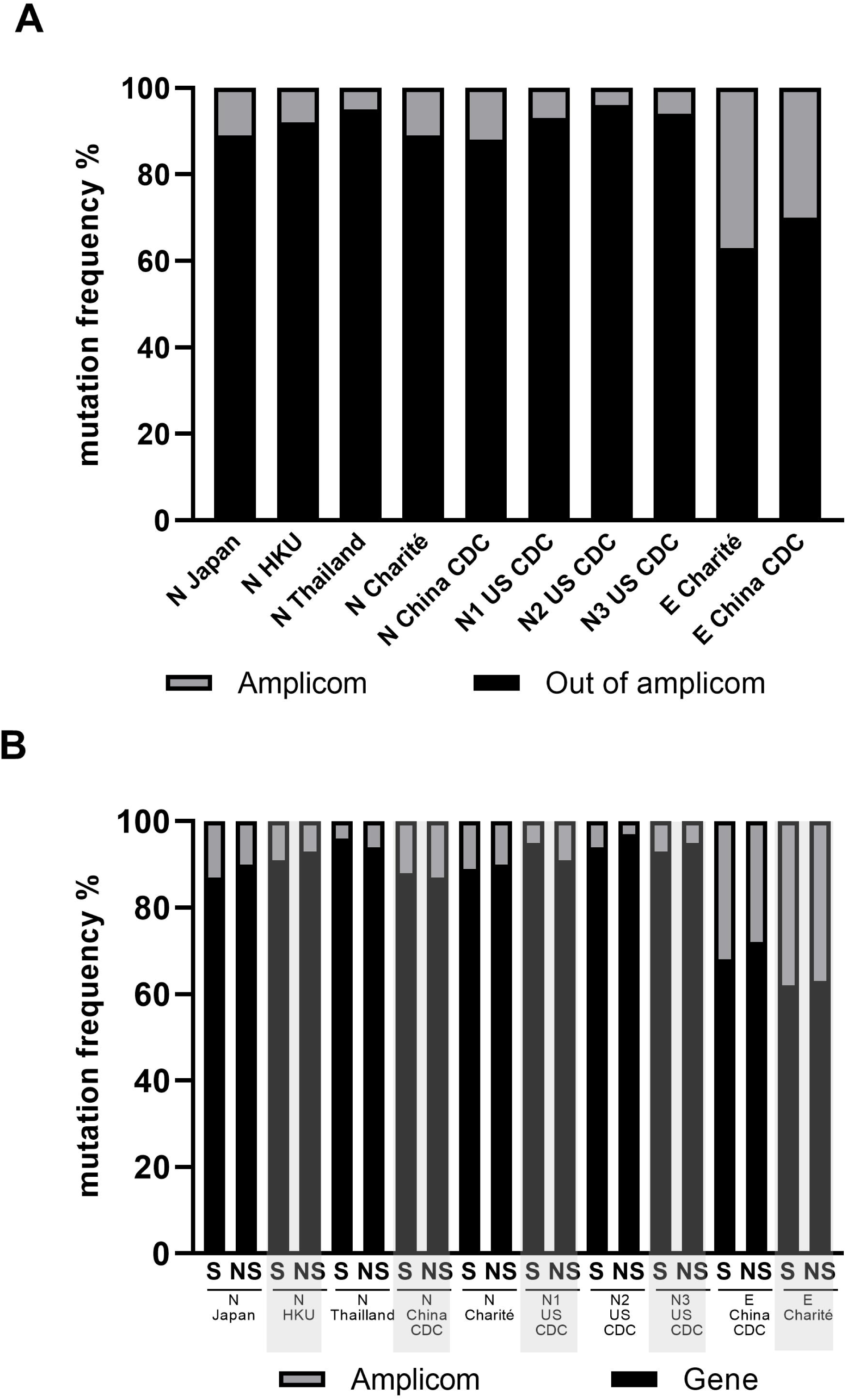
Mutation frequencies in E and N targets to identification of SARS-CoV-2. Frequency of mutations in gene targets and amplicon used by RT-qPCR protocols were quantified by Target mutations® software as described in methods. Data on the graphs represent the frequency mutations in (A) whole gene or amplification regions and (B) discriminated data by (S) synonymous and (NS) non-synonymous mutations. A total of 1512 and 185 mutations were identified in the entire NP and E gene, respectively, as of February 27, 2023. CDC: Centers for Disease Control and Prevention; HKU: Hong Kong.

### Identifying mutation that compromise RT-qPCR protocol in a world scale

Since many mutations are reported in the regions of SARS-CoV-2 genome used in the RT-qPCR protocols to molecular identification [5], we ask if this SNPs can impair the protocol capacity to identify the new variants of the virus. We stablished a workflow to validate the RT-qPCR primer and probes sets by the sorting of the new variant representative genomes around the world using GSAID initiative (Figure 2A). We identified just four fixed mutations in the Omicron variant in the regions of the molecular protocols (Table 2). In this sense, SNPs present in the N target of the CDC China indicates as problem of the primer design [12], since all SARS-CoV-2 the mutations including the reference strain (Table 2). Another problem in the primer design was identified in RdRp protocol from CDC China, since primer and probe do not annealed with targets in all SARS-CoV-2 variants [12]. Three fixed SNPs were identified in the amplicon regions of the Omicron variant: (i) c.38C/T int TaqMan probe of N1 US CDC protocol; (ii) c.26C/T in the forward primer of E Charité protocol (Figure 2D) and; (iii) c.2010A/G in the forward primer of R Charité protocol (Figure 2C). Deletion of nine nucleotides (c.del89_98) in the location of reverse primer indicates that the NIH Thailand protocol is not able to identify Omicron variants (Figure 2D and Table 2). Then, the data suggest few SNPs are conserved in new variants around the world and just a deletion in the region recognized by one RT-qPCR primers and probe set was compromised.

**Figure 2.**
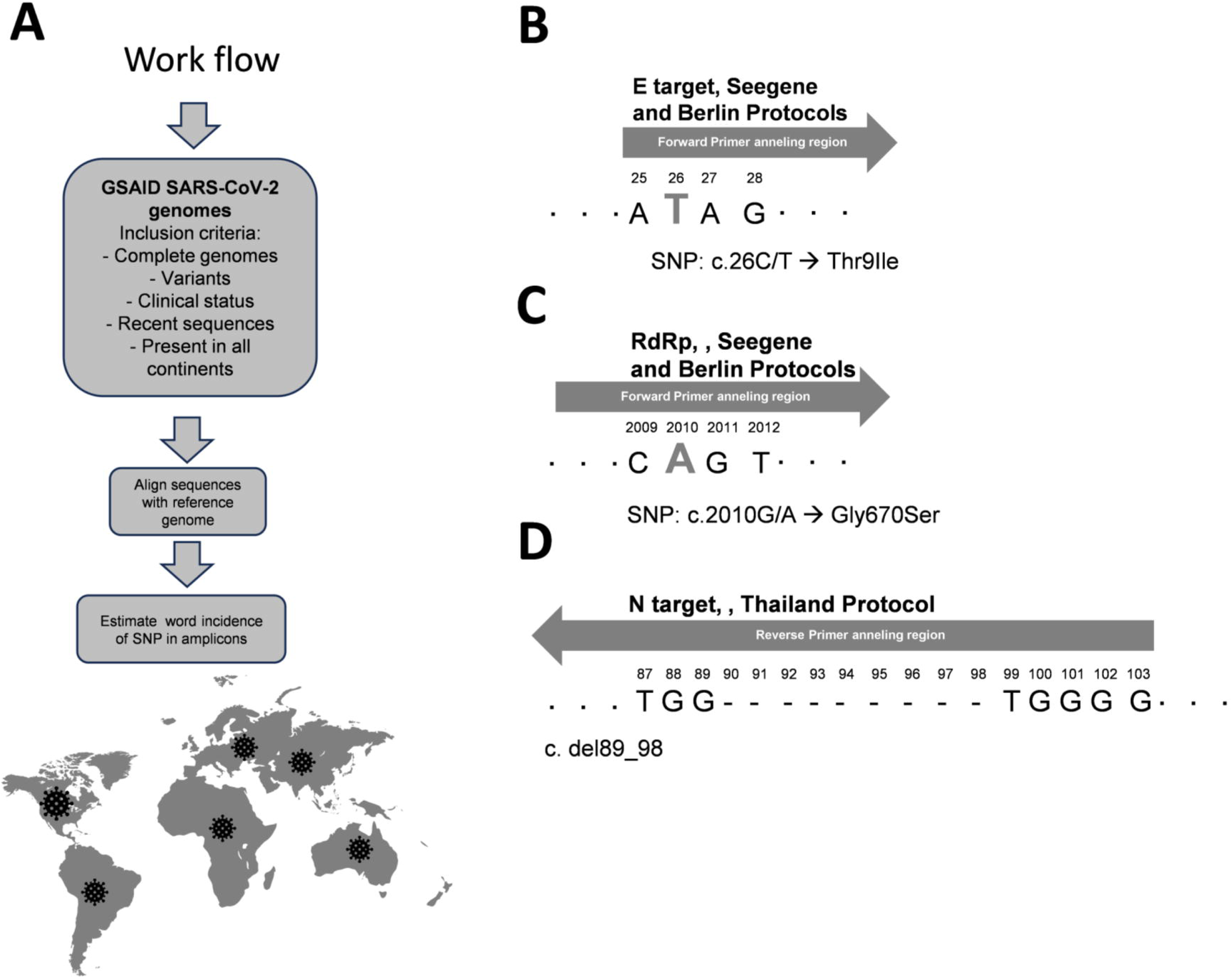
Identifying the prevalent mutations in a world scale in the primer annealing regions of the RT-qPCR protocol used for detecting SARS-CoV-2. (A) The workflow to identify prevalent mutations was stablished by recovering genome sequences in the GSAID data bank as described in methods. Recent and random samples were collected using quality criteria. Diagrams represent mutations in regions of annealing primers compromised for SARS-CoV-2 detection. Single nucleotide polymorphisms (SNP) were detected in the forward primer annealing region of the (B) Envelop protein and (C) RNA polymerase dependent of RNA (RdRp) genes. (D) Deletion of nine nucleotides were identified in the nucleocapsid phosphoprotein gene in the reverse primer annealing region.

**Table 2.**
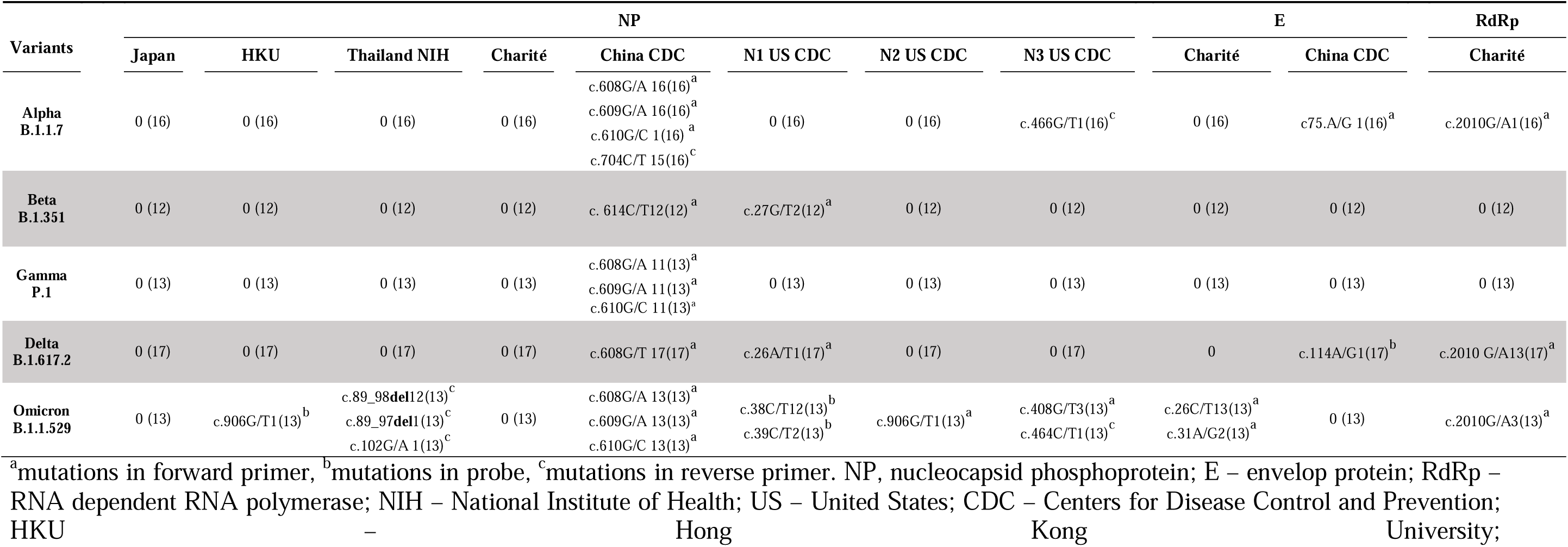
Single nucleotide polymorphisms in the haplotypes of SARS-CoV-2 variants derived from different continents.

### SNPs in the primer and probe sets do not affect detection of the new variants

Frequency of mutations in the amplicon regions of the RT-qPCR protocols are used to justify invalidating of several methods [5]. Many articles use few samples of new SARS-CoV-2 variants to identify sensibility and specificity of the molecular tests [1,13–15]. We evaluated the effect of SNPs in the efficiency of the protocols to detect SARS-CoV-2 new variants. To this, we design long oligos as template to RT-qPCR protocols containing SNPs identified in the Omicron variants regions of the primer annealing to the RdRp and E genes of the Charité and Seegene protocol. We observed that both protocols can detect variants containing fixed SNPs in the E and RdRp gene, with a detection limit of ten molecules per reaction (Figure 3). Then, the data suggest that haplotypes containing a SNP continue to be recognized by primers and probes designed to the SARS-CoV-2 reference genome.

**Figure 3.**
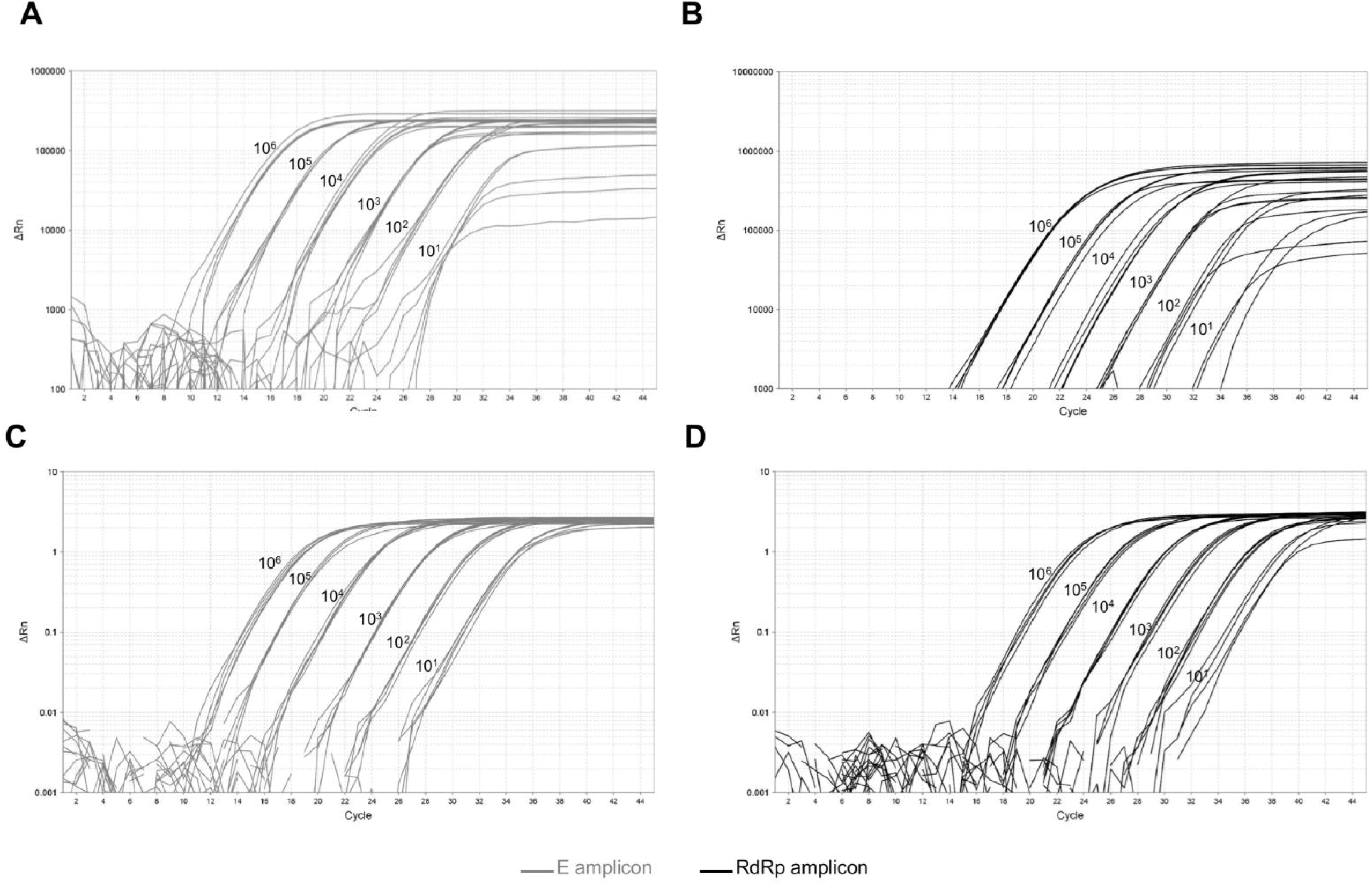
Checking the primer detection of prevalent SNPs in E and RdRp targets of news variants of SARS-CoV-2. Long oligos were designed to mimicry SNPs found in new variants in regions of the annealing primers with E gene c.26C>T or RdRp gene c.2010G/A. Logarithmic amplification plots represent RT-qPCR test using serial dilution of synthetic E (grey lines) and RdRp (black lines) muted long oligos using (A-B) Seegene reagent kit or (C-D) Berlin protocol (Corman et al. 2020).

## DISCUSSION

The ability of molecular diagnostic tests based on the RT-qPCR technique to detect new variants of SARS-CoV-2 has been investigated [1,5,15]. Some work has relied on single base mutation frequencies in databases containing a few thousand sequences to determine which tests should be discarded [5]. Herein, we established a workflow using data available in genome databases to investigate whether the primer and probe sets used in the first wave of the COVID-19 pandemic are still for new variants, and we verified that most molecular tests for SARS-CoV-2 detection during first wave of COVID-19 continue detecting new variants of the virus.

The COVID-19 pandemic tested the limits of the RT-qPCR technique. Previously restricted to a few pathologies and present in few laboratories in Brazil and in many countries, the application of the technique needed to be expanded to different locations, which allowed the establishment of the sensitivity and specificity of different RT-qPCR protocols for the identification of SARS-CoV-2 in biological samples [1,15]. Likewise, genetic sequencing has been widely carried out, with more than 3 million viral genomes currently available in genetic databases [8].

Although the RT-qPCR test is considered the gold standard for diagnosing SARS-CoV-2, patients who were negative for the test when submitted to chest computed tomography exams presented typical findings of the disease, such as ground-glass opacity. The same patients subjected to isolation, after repeated swab tests, received a positive result, indicating that depending on the patient’s clinical stage, RT-qPCR tests may present false-negative results [16,17]. False negative tests by RT-qPCR can be explained by the presence of mutations in the annealing regions of primers and probes. Some studies that tested the sensitivity of the RT-qPCR protocols available for diagnosing SARS-CoV-2 found that annealing regions of the set of primers and probes from protocol N from Japan and ORF1ab from CDC China presented mutations, however, it did not affect the sensitivity of these protocols [18]. Here, it was also identified that the protocols remain valid, but without the need to resort to biological samples for validation.

Regarding the mutations found in the SARS-CoV-2 genome and their influence on RT-qPCR diagnostic tests, it has been reported that the loss of six nucleotides in the S gene (207_212del) is associated with errors in the results of tests that use TaqPath probe targeting the S gene [19,20].

A recurrent SNP at position 26340 of the virus genome, which it inside of the E gene, was reported to be responsible for making a commercial test, Roche’s COBAS test, unfeasible [21]. Despite, other molecular tests using the same primer and probes of COBAS test are not affected [21]. Here, it was definitively identified that this mutation does not alter the efficiency of the reaction in the protocols used which have position 26340 as reference, among those Charité protocol [10] or the Allplex protocol from the Seegene company. Different haplotypes in the B.1.1.7 lineage was identified, but those mutations are not sufficient to invalidate the target of the tested Berlin-Charité protocol [22]. In another work, in which they investigate targets within South America, the authors indicate that the E gene is the most conserved [23]. However, these authors should consider the size of the genes used in their analyses. Here, a lower relative frequency of mutations was identified in the E gene compared to the N gene, but it must be considered that the E gene has five times fewer nucleotides than the N gene.

To try to solve the problem of the impact of variants on RT-qPCR tests, other work indicates the use of multiplex assays, in which the combination of two genes would make the result more robust and thus less susceptible to interference from mutations [24]. Furthermore, screening tests have also been proposed to previously identify SARS-CoV-2 variants in samples, however limitations have been pointed out for cases in which there are few deletions/mutations in these diagnoses [25].

Currently, so-called Virus-Like Particles or VLPS are being proposed to study variants because they allow the assessment of mutations in all structural proteins [26]. VLPS could be used as an alternative for validating diagnostic methods. However, regarding molecular diagnosis, a more economical alternative is the use of polynucleotides containing the most prevalent SNPs in circulating variants, as was conducted in the present work. From this approach it was identified that a single polymorphism in primer annealing regions did not reduce their detection capacity in samples. The combination of genome analysis of new variants with the possibility of in silico design of sequences that mimic SNPs identified by bioinformatics can facilitate monitoring the effectiveness of already established tests in an easier and faster way and with a lower risk for researchers. This strategy can be used to monitor the effectiveness of tests against emerging viruses in new pandemic situations. Furthermore, similar strategies can be applied to determine the validity of existing tests against viruses from the Influenza family, for example. Polynucleotides can also be designed to mimetize the sequence of variants, reducing the need for empirical testing using isolated viruses.

This study has a potential limitation, since SNPs for RdRp gene is poor investigated in the data bank used. Then, we decide to do not include the data of RdRp in the SNP analysis in this study (https://www.ncbi.nlm.nih.gov/labs/virus/vssi/#/scov2_snp). Deposit of more haplotypes including RdRp gene SNPs are necessary to better understand the impact of SARS-CoV-2 evolution in the RT-qPCR diagnosis using primers and probes that annealing in this gene.

In sum, our data suggest that majority of the molecular targets designed for the first SARS-CoV-2 variants remain valid for the identification of new virus variants despite the large number of identified haplotypes. However, false negative test failures can be identified by using more than one molecular target for the same sample. Regions that are under viral selective pressure should be avoided in the use of the diagnostic, once new variants are selected, molecular tests are affected on a global scale.

## Author Contributions

Conceived and designed the experiments: AFRA, RCK and TA-S. Data collection: AFRA, WAA, WMP, TA-S; Analyzed the data: AFRA, WAA, WMP, RSS, LPRV, RCK, MHFK, JBL, TA-S; Contributed materials/analysis tools: AFRA, WAA, WMP,RSS, TA-S; Wrote the paper: AFRA, TA-S.

